# MorphoLearn: A morphology-driven workflow to decipher 3D electron microscopy segmentation in diatoms

**DOI:** 10.64898/2026.01.15.699631

**Authors:** Clarisse Uwizeye, Serena Flori, Jhoanell Angulo, Pierre-Henri Jouneau, Benoit Gallet, Pascal Albanese, Giovanni Finazzi

## Abstract

Three-dimensional electron microscopy (3D EM) enables the quantitative analysis of cellular ultrastructure. However, large-scale segmentation of whole-cell volumes poses a significant challenge, especially in biologically diverse systems. Unlike medical and animal cell imaging, which often benefit from temporal redundancy and relatively uniform morphology, studies of microbial and microalgal biodiversity must rely on static snapshots. These snapshots exhibit high variability in cell shape, organelle organisation, and image contrast. Consequently, robust AI-assisted segmentation in this context requires models that learn directly from morphological features and can adapt to heterogeneous sample preparation. In this paper, we present a systematic framework for AI-assisted segmentation of Focused Ion Beam-Scanning Electron Microscopy (FIB-SEM) datasets. This framework is specifically designed to address the challenges posed by morphological diversity and contrast variability while remaining within realistic computational constraints. We evaluate multiple lightweight 3D encoder-decoder architectures and identify VNet as the best option for balancing computational efficiency and volumetric accuracy in whole-cell segmentation. Using datasets from two strains of *Phaeodactylum tricornutum* and extending our analysis to cross-species comparisons, we demonstrate that training on specific regions of interest can lead to an overestimation of model performance. In contrast, performing whole-cell segmentation uncovers significant differences in architectural robustness. Moreover, we show that transfer learning and contrast-aware hybrid strategies allow for efficient adaptation to previously unseen datasets with minimal annotation. The incorporation of boundary-aware loss functions significantly enhances the delineation of closely associated organelles, such as chloroplasts and mitochondria, in multi-class segmentation tasks. Together, these findings establish a scalable, reproducible, and biologically informed AI framework for 3D FIB-SEM segmentation. This framework enables high-throughput analysis of cellular ultrastructure across diverse species and imaging conditions.

**Author Summary:** Cells exhibit a wide range of shapes, sizes, and internal structures, particularly among various microbial species. These morphological differences are not arbitrary; they indicate how cells adapt to their environments and manage essential biological functions. Modern three-dimensional electron microscopy can capture this structural diversity at the nanometre scale, but analysing the resulting data is often slow. This is due to the time-consuming process of manually outlining cellular structures, which also requires expert knowledge. Artificial intelligence (AI) has made significant advances in accelerating image analysis in medical and animal cell studies, typically by learning from repeated observations over time. However, studies focusing on microbial and microalgal biodiversity often rely on single snapshots of diverse cells prepared under varying imaging conditions. This complicates automated analysis since AI systems must learn from morphology directly rather than from temporal repetition. In this study, we developed and evaluated an AI-assisted segmentation framework specifically for whole-cell 3D electron microscopy data. By systematically comparing efficient neural network architectures and incorporating transfer learning and contrast-aware strategies, we demonstrate that accurate segmentation can be achieved even with limited training data and standard computing resources. Our approach facilitates faster, scalable, and reproducible analysis of cellular ultrastructure, paving the way for large-scale investigations into cell morphology, adaptation, and diversity across species.

**Significance statement:** Quantitative analysis of cellular ultrastructure across species is currently limited by challenges in segmenting large three-dimensional electron microscopy datasets. Unlike medical imaging, which often benefits from artificial intelligence due to its use of temporal repetition and consistent morphology, studies of microbial biodiversity depend on single snapshots that display extreme variations in cell shape and image contrast. This work presents a scalable, morphology-driven AI framework for whole-cell 3D segmentation that is resilient to biological diversity and variations in sample preparation. By enabling accurate analysis with minimal annotations and standard computational resources, this approach enhances access to high-throughput ultrastructural studies and facilitates comparative investigations of cellular adaptation across different species.

## Introduction

Cell morphology originates from a tightly regulated spatial organisation of subcellular compartments that supports cellular function and adaptation. In unicellular organisms, diatom morphology is highly plastic; cell size, shape, and organelle arrangement can change rapidly in response to environmental conditions, nutrient availability, or physiological stress. These morphological adaptations provide rich biological information, reflecting the cellular state, acclimation strategies, and survival mechanisms. Therefore, understanding how morphology varies across different cells, conditions, and species is central to studying cellular diversity and adaptation.

In many biomedical and animal cell systems, artificial intelligence (AI) approaches benefit from temporal sameness. Time-lapse imaging allows models to learn from repeated observations of the same cell or lineage, where morphology evolves gradually, and differences remain relatively stable. In contrast, studies of microbial and microalgal biodiversity face a fundamentally different challenge: cells are often imaged as static snapshots across species with distinct morphologies and under variable preparation and staining conditions. In this context, AI must learn directly from morphological structures instead of relying on temporal continuity. Effective AI-assisted segmentation in biodiversity studies requires models that can generalise across significant variations in shape, size, organelle organisation, and image contrast.

Recent advances in three-dimensional electron microscopy (3D EM) have enabled unprecedented access to cellular ultrastructure (McCafferty et al, 2024). Cryo-electron tomography (cryo-ET) has been transformative for resolving subcellular architecture in near-native states, providing detailed views of macromolecular organisation (Lamm et al., 2024; Yamauchi et al., 2024). However, cryo-ET remains limited in throughput and typically focuses on small subcellular regions. Preparing and milling cryo-lamellae for whole-cell reconstruction is time-consuming and often requires several days per cell, making large-scale morphological comparisons impractical.

Focused Ion Beam-Scanning Electron Microscopy (FIB-SEM) offers a complementary approach by enabling volumetric imaging of entire cells or multiple cells, depending on their size, within a single acquisition. Although FIB-SEM does not preserve native hydration, it offers continuous 3D information at voxel resolutions of 5-10 nm, making it particularly suitable for studying cellular variability and morphological plasticity across populations. Importantly, FIB-SEM allows for the acquisition of dozens of whole-cell volumes within hours rather than days, enabling statistically meaningful analyses across different conditions and species(Uwizeye et al., 2021; Uwizeye et al.,2021; Ezzedine et al., 2023; Catacora Grundy et al., 2023; Permann et al., 2023). However, this advantage is offset by a significant bottleneck: manual segmentation of FIB-SEM datasets typically requires several days of expert annotation per cell, severely limiting scalability.

Focused Ion Beam-Scanning Electron Microscopy (FIB-SEM) offers a complementary approach by enabling volumetric imaging of entire cells or multiple cells within a single acquisition. Although FIB-SEM does not preserve native hydration, it offers continuous 3D information at voxel resolutions of 5-10 nm, making it particularly suitable for studying cellular variability and morphological plasticity across populations. Importantly, FIB-SEM allows for the acquisition of dozens of whole-cell volumes within hours rather than days, enabling statistically meaningful analyses across different conditions and species. However, this advantage is diminished by a significant offset: manual segmentation of FIB-SEM datasets typically requires several days of expert annotation per cell, severely limiting scalability.

AI-assisted segmentation has the potential to overcome this limitation, yet most deep learning approaches have been developed and validated in medical imaging contexts (Ekman et al., 2025). Many architectures were initially designed for 2D data and rely on relatively homogeneous morphology and consistent contrast (Wang et al., 2022; Singh et al., 2020). While such models can perform well with limited data, they often struggle to capture the volumetric context and structural complexity that define subcellular organisation in 3D EM datasets (Fauser et al., 2019; Ding et al., 2023; Faghani et al., 2023). Extending these architectures to three dimensions substantially increases computational and memory requirements, making efficient, lightweight implementations essential for practical use with large volumetric datasets.

Recent studies have shown that carefully designed lightweight 3D architectures can achieve performance comparable to larger models on small datasets, suggesting that efficiency and accuracy are not mutually exclusive (Perera et al., 2024; Huang et al., 2019). However, most existing work focuses on single species, uniform contrast conditions, or narrowly defined segmentation tasks. *Two critical challenges remain largely unexplored concerning FIB-SEM segmentation: generalisation across biologically diverse cell morphologies and robustness to contrast variability arising from differences in sample preparation and staining*.

In this study, we present a proof-of-concept 3D deep learning framework designed specifically to address these challenges. Rather than relying on temporal information, our approach learns directly from morphology, enabling segmentation across diverse microalgal species and variable imaging conditions. We systematically benchmark lightweight 3D encoder-decoder architectures under realistic computational constraints and integrate transfer learning, contrast-aware training, and hybrid designs to enhance robustness and scalability. By explicitly targeting morphological diversity and contrast heterogeneity, our framework provides a practical and reproducible solution for large-scale 3D segmentation of FIB-SEM datasets, enabling quantitative ultrastructural analysis across different cells, conditions, and species.

## Materials and methods

### Datasets

We analysed three-dimensional Focused Ion Beam-Scanning Electron Microscopy (FIB-SEM) datasets to characterise the ultrastructure of the diatom *Phaeodactylum tricornutum* (strains 1 and 18). We aimed to compare intraspecific features and exploit *Thalassiosira* data for a cross-species comparison (see Figure 1). Including a second species was intended to test the generalizability of our model beyond a single species. All datasets consisted of isotropic voxel volumes with resolutions ranging from 5 to 10 nm per pixel, allowing us to reconstruct entire cells and identify both simple subcellular compartments (e.g., vacuoles) and more complex organelles (e.g., chloroplasts and mitochondria). Before analysis, we preprocessed the raw volumes to reduce noise using median filtering. The processed volumes were then subdivided into fixed-size sub-volumes for model training and inference. Ground-truth annotations were manually generated in 3D Slicer to create binary segmentation masks for each organelle of interest. As a proof of concept, we initially adopted a one-class segmentation strategy (foreground versus background) to assess the model’s ability to learn structural features from limited annotated data. We then extended this approach to multi-class segmentation (Table 1).

**Figure 1:**
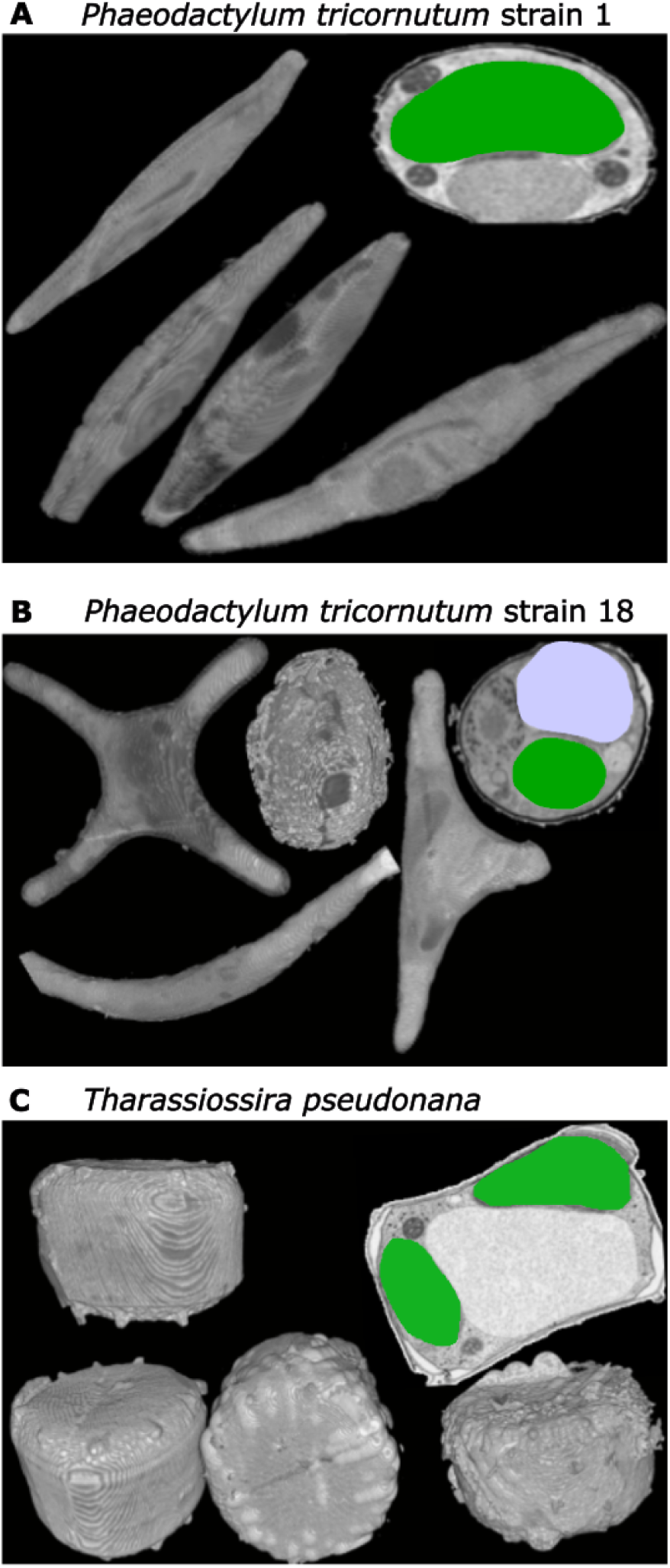
Representative 3D FIB-SEM reconstructions of the model diatoms used in this study. (A) *Phaeodactylum tricornutum* strain 1 is characterised by its elongated, fusiform shape and distinct intracellular organisation, which includes chloroplasts, lipid droplets, mitochondria, and a nucleus. (B) *Phaeodactylum tricornutum* strain 18 exhibits a high degree of morphological plasticity, with cells ranging from cruciform to triradiate, fusiform, and oval, and with vacuoles. (C) *Thalassiosira pseudonana* displays compact, cylindrical frustules with well-defined chloroplast organisation and a large vacuole. In the diagrams, the interesting organelles are labelled in green for chloroplasts and light purple for vacuoles. These samples illustrate the morphological diversity and subcellular complexity captured in FIB-SEM datasets, forming the basis for AI-assisted segmentation and model benchmarking. The complete training dataset is available on **BioImage Archive** (S-BIAD2407).

**Table 1:** Computing resources and dataset specifications

| Resource | Details |
| --- | --- |
| Operating system (OS) | Linux and windows |
| Computing infrastructure | GPU computer, GPU cluster |
| GPU capacity | PC windows: RTX A2000: 12GB; (Gricad cluster): A100: 32-64 GB |
| Training datasets | 32 stacks and 32 labels (ground truths) |
| <u>Additional data</u> | <u>FIB-SEM data to evaluate</u> |
| **Sample<sub>1</sub>:**<br>*Phaeodactylum tricornutum* 1 | 42 stacks |
| **Sample<sub>2</sub>:**<br>*Phaeodactylum tricornutum* 18 | 39 stacks |
| **Sample<sub>3</sub>:**<br>*Thalassiosira pseudonana* | 12 stacks |

### Data augmentation

To address the challenges posed by the limited size and diversity of FIB-SEM datasets, we implemented data augmentation techniques to improve model generalisation and reduce overfitting on the training data. The augmentations were applied only to the training set. The following transformations were used: (i) random flipping along all spatial axes (x, y, z); (ii) the addition of Gaussian noise to simulate artifacts commonly found in electron microscopy; (iii) random contrast adjustments to account for variations in staining; (iv) affine transformations to introduce controlled spatial distortions; and (v) zoom operations to accommodate differences in organelle size. All transformations were explicitly designed to maintain organelle topology and structural integrity (see Figure 2). Data augmentation was performed on-the-fly during training, generating unique image variants during each iteration. This strategy effectively increased the size of the training dataset by over an order of magnitude and enhanced the model’s robustness to previously unseen samples. Furthermore, when combined with transfer learning, data augmentation was crucial to our methodology, demonstrating that accurate segmentation can be achieved even with limited annotated data (Table 2).

**Figure 2.**
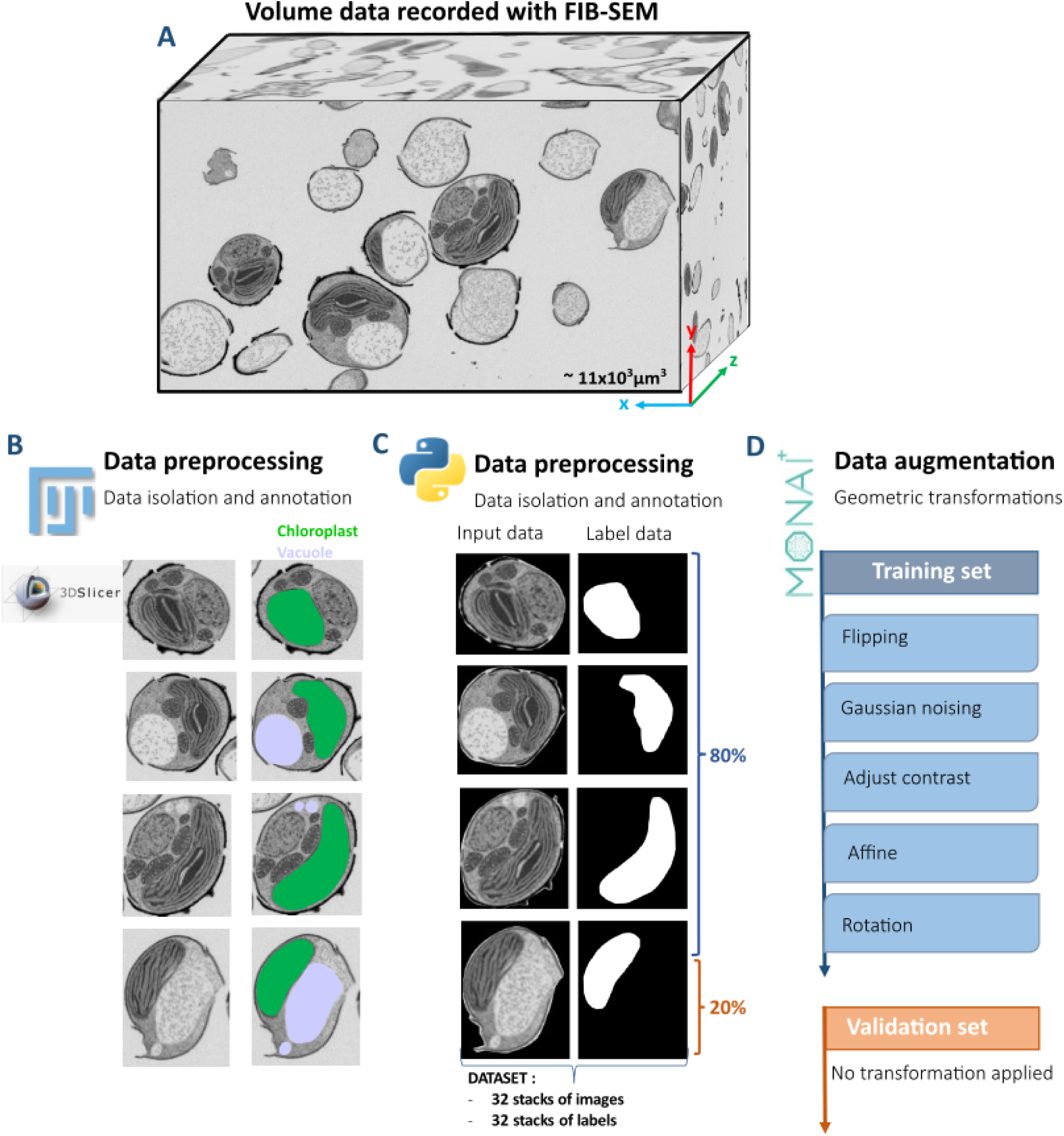
Workflow for Processing Volumetric Diatom Images and Preparing Datasets. (A) Example of a three-dimensional volume dataset obtained through focused ion beam–scanning electron microscopy (FIB-SEM), which reveals cellular ultrastructure in high-resolution voxel space. (B) Data preprocessing and manual annotation of organelles, including chloroplasts (green) and vacuoles (violet), from volumetric image stacks. (C) Data collection focused on preparing paired input and label data, converting them into NIfTI format for model training.. (D) A data augmentation pipeline is implemented using MONAI, incorporating geometric and photometric transformations such as flipping, addition of Gaussian noise, contrast adjustment, affine transformations, and rotation. The dataset is divided into training (80%) and validation (20%) sets, with augmentations applied only to the training data.

**Table 2:**
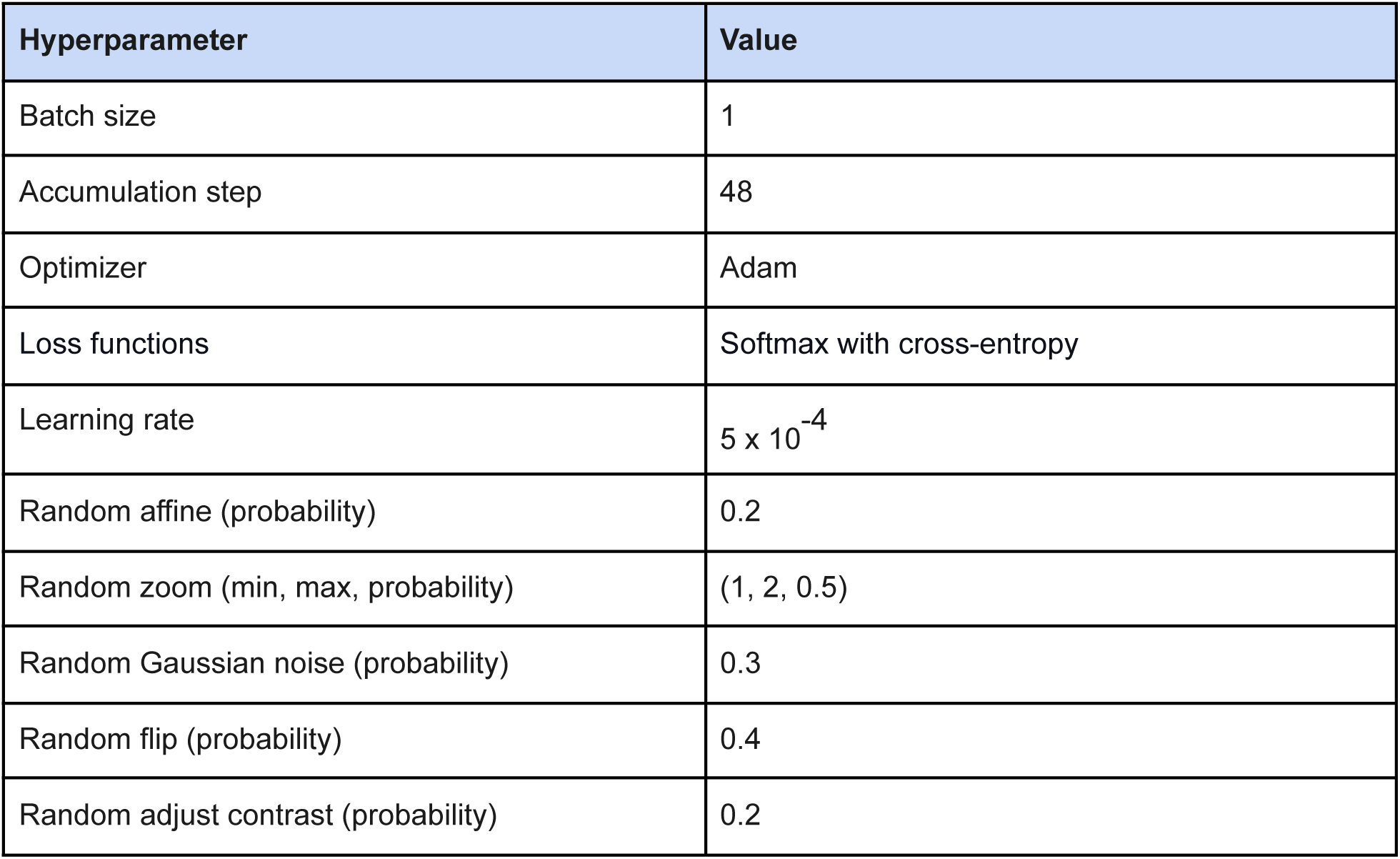
Model hyperparameters for object segmentation

| Hyperparameter | Value |
| --- | --- |
| Batch size | 1 |
| Accumulation step | 48 |
| Optimizer | Adam |
| Loss functions | Softmax with cross-entropy |
| Learning rate | $5 \times 10^{-4}$ |
| Random affine (probability) | 0.2 |
| Random zoom (min, max, probability) | (1, 2, 0.5) |
| Random Gaussian noise (probability) | 0.3 |
| Random flip (probability) | 0.4 |
| Random adjust contrast (probability) | 0.2 |

#### Hyperparameter settings

### Network architecture design: evaluation of four-level architectures

To identify an effective and computationally efficient architecture for 3D FIB-SEM segmentation, we implemented and compared three four-level convolutional neural network (CNN) encoder-decoder architectures commonly used for volumetric image analysis (see Fig. 3). All models were adapted for 3D inputs and designed to operate within moderate computational constraints (≤ 12 GB GPU memory).

**Figure 3.**
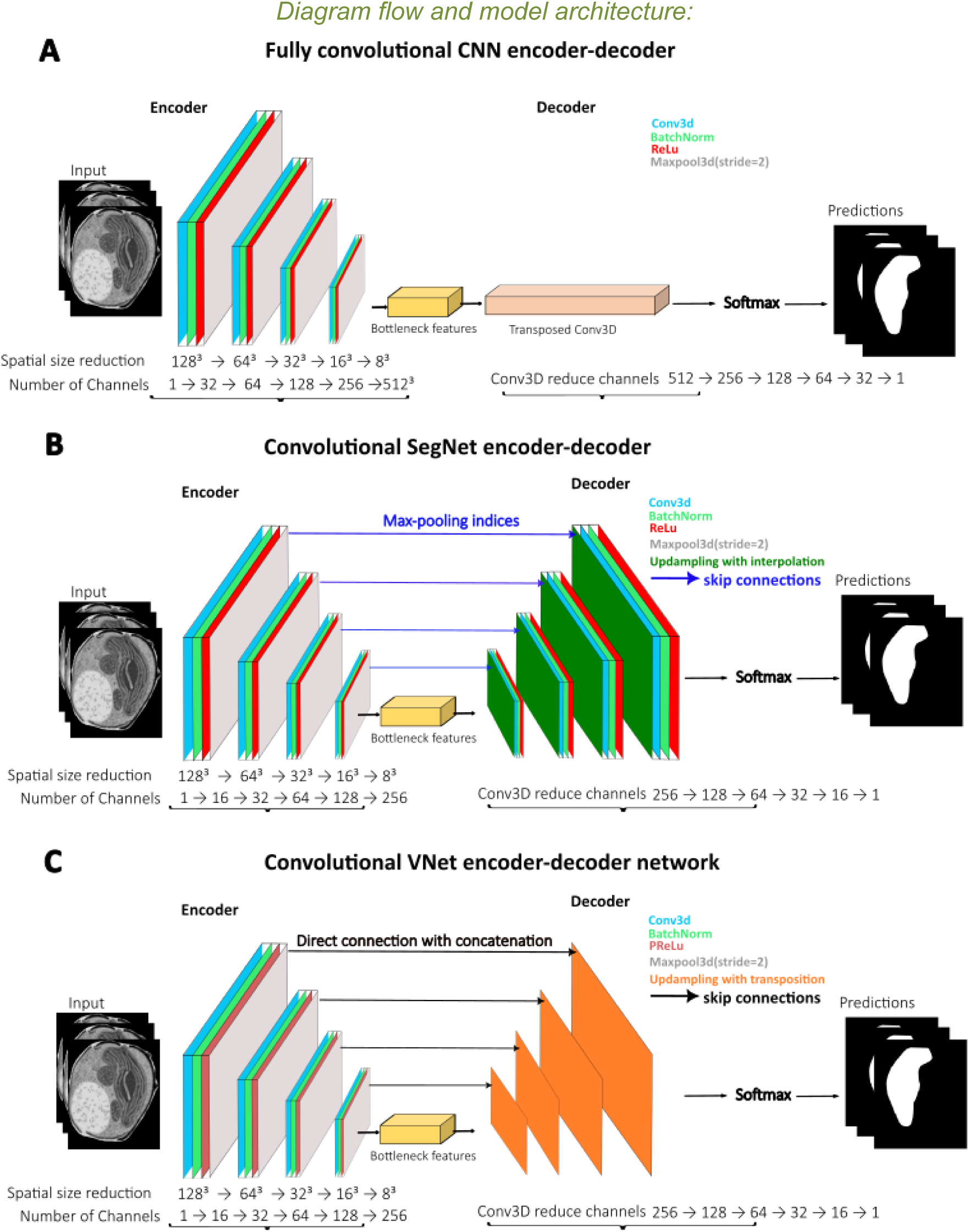
Comparison of isolated 3D convolutional encoder-decoder architectures used for proof-of-concept evaluation. (A) A fully convolutional **CNN** encoder-decoder featuring symmetric encoding and decoding blocks. (B) The **SegNet** architecture, which utilizes max-pooling indices and skip connections to achieve spatial reconstruction. (C) The **VNet** architecture, which employs direct concatenation between the encoder and decoder to maintain volumetric context. Each model processes 3D FIB-SEM sub-volumes to predict binary segmentation masks. See details in supplementary information (**STable 1** and **STable 2**)

Figure 3A illustrates a fully convolutional 3DCNN encoder-decoder architecture. This model progressively downsamples spatial resolution through a series of stacked convolutional and pooling layers to extract hierarchical features. It is followed by a symmetric decoder that aims to restore spatial resolution using transposed convolutions. This architecture serves as a minimal baseline, lacking explicit skip connections or pooling index reuse.

Figure 3B shows a SegNet-style encoder-decoder (3DSegNet), where max-pooling indices calculated during the encoding phase are reused during decoding to guide upsampling. This design includes skip connections between corresponding encoder and decoder levels, enhancing the recovery of spatial details while maintaining computational efficiency.

Figure 3C presents a VNet-based architecture (3DVNet) that features direct skip connections between the encoder and decoder stages through feature map concatenation. This design preserves fine-scale structural information and improves boundary delineation, making it particularly suitable for segmenting complex 3D ultrastructural features.

All three architectures utilised the same foundational 3D convolutional blocks at each level. These blocks consisted of Conv3D layers followed by batch normalisation, ReLU or PReLU activation, and max pooling. This controlled design ensured that any performance differences primarily reflected the architectural topology, rather than variations in parameter count or training strategy. These baseline architectures formed the reference models evaluated in Phase 1. Subsequent benchmarking against established medical imaging frameworks and hybrid extensions is discussed in Phases 2 and 3.

#### Phase 2-Benchmarking Against Established 3D Medical Imaging Frameworks

In this phase, we conducted a performance benchmark of the selected baseline architecture against established 3D segmentation frameworks under identical training conditions. Based on the results from Phase 1, we chose the VNet-based architecture (3DVNet; see Figure 3C) as the reference model due to its superior ability to preserve fine structural details via direct encoder-decoder skip connections. This architecture was then benchmarked against widely used 3D medical imaging frameworks implemented in MONAI, including BasicUNet and DynUNet.

All models were trained and evaluated under the same conditions to ensure a fair comparison. This included using identical input volumes, augmentation strategies, optimisation settings, and computational constraints (limited to ≤ 12 GB of GPU memory). To assess model stability and generalisation in scenarios with limited annotated data, we employed a three-fold cross-validation protocol. Additionally, we evaluated Swin-Net, a transformer-based model originally developed for 2D ultrasound image segmentation that has recently demonstrated high segmentation accuracy (Zhu et al., 2024). We adapted Swin-Net to support 3D volumetric inputs, allowing for direct comparison with convolutional models. This evaluation enabled us to determine whether incorporating broader spatial context through self-attention mechanisms can enhance segmentation performance for 3D electron microscopy data.

#### Phase 3 - Hybrid architectures and contrast-aware learning

In this final phase, we explored hybrid network designs that combine the strengths of convolutional and transformer-based architectures, optimising transfer learning to improve data efficiency and robustness to imaging variability.

The first hybrid configuration integrated Swin-Net and 3DVNet into a unified architecture called 3DSwin-VNet. This model merged the global contextual modelling capabilities of the transformer-based Swin encoder with the strong local feature continuity of VNet. It employed parallel encoding pathways followed by a shared decoding stage. Both encoders were initialised with pre-trained weights from earlier training phases. This approach allowed us to retain previously learned structural representations and accelerate convergence. We tested whether jointly modelling global context and local continuity improved segmentation consistency in complex 3D electron microscopy data.

The second hybrid configuration implemented a contrast-aware dual-encoder 3DVNet. In this setup, separate encoders were trained on high- and low-contrast image subsets to capture contrast-specific intensity profiles. These latent feature representations were then merged in a shared decoder. This contrast-specific training strategy aimed to reduce sensitivity to staining variability and improve generalisation across datasets acquired under different preparation protocols or imaging settings.

For both hybrid designs, we fine-tuned the models using reduced learning rates (ranging from 1×10^-5^ to 5×10^-5^), small batch sizes, and gradient accumulation. This helped preserve pre-trained features while allowing gradual adaptation to the target domain. Training was concluded when validation loss converged, or early stopping criteria were met. All hybrid models were implemented in lightweight configurations to ensure compatibility with standard GPU resources. This supports our overall goal of developing an accessible, scalable, and reproducible segmentation framework for 3D electron microscopy.

#### Training, validation, and optimisation

All models were trained using supervised voxel-wise segmentation, employing binary cross-entropy (BCE) as the primary loss function for single-class experiments. BCE was selected for its computational efficiency and stability when dealing with large 3D volumes. Model optimization was carried out using the Adam optimiser, with a learning rate of 1×10^-4^, β_1_ = 0.9, and β_2_ = 0.999.

To manage limited GPU memory while maintaining training stability comparable to larger batch sizes, we used gradient accumulation, with up to 48 accumulation steps. The models were trained for 300-500 epochs, with early stopping triggered if the validation loss did not improve for more than 20 consecutive epochs. To evaluate robustness and generalisation, all experiments were performed using 3-fold cross-validation, with each fold used once as the validation set. For benchmarking experiments involving BasicUNet and DynUNet, we adhered to the three-fold cross-validation protocol recommended by MONAI for small datasets. Validation was conducted on held-out volumes that were not used during training or fine-tuning. Data augmentation techniques were applied only during the training phase, as described previously.

#### Fine-tuning strategy

To address differences in structural complexity across organelles, we implemented a two-stage fine-tuning protocol. For organelles with simple morphology, the models were first trained on cropped regions of interest (ROIs) to capture local features, followed by fine-tuning on complete volumetric data to incorporate global context. Conversely, for structurally complex organelles, models were initially trained on complete volumes to learn contextual organisation, followed by ROI-based fine-tuning to enhance boundary precision. This strategy allowed for efficient learning while reducing annotation requirements.

#### Performance metrics

Segmentation performance was evaluated using three complementary metrics: Binary Cross-Entropy (BCE), Dice coefficient (D), and F1-score (F_1_). BCE quantified voxel-wise probabilistic accuracy, while Dice and F_1_ assessed spatial overlap between predicted segmentations and ground-truth labels, providing robustness to class imbalance.

The Dice coefficient was defined as:

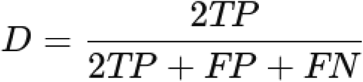

Where TP, FP, and FN denote true positive, false positive, and false negative voxels, respectively.

The F1-score is equivalent to the Dice coefficient for binary segmentation tasks:

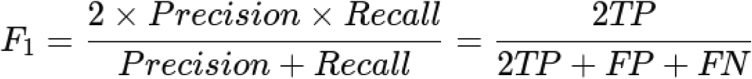

Together, these metrics provided a balanced evaluation of voxel-level accuracy and volumetric reconstruction quality across baseline, hybrid, and transfer-learned models.

#### Loss Functions

Model optimization employed a composite loss combining Binary Cross-Entropy and Dice loss for single-class segmentation experiments, balancing voxel-wise accuracy with regional overlap. This formulation was chosen for its low memory footprint and stable convergence properties, making it suitable for benchmarking multiple 3D architectures under realistic computational constraints (Milletari et al., 2016; Sudre et al.,2017, Ma et al., 2021). For multi-class segmentation experiments, an additional Hausdorff Distance (HD) loss term was incorporated to improve boundary accuracy and inter-class separation. The Hausdorff distance loss penalises large deviations between predicted and ground-truth object surfaces and was implemented in the MONAI framework, which computes a differentiable 3D Euclidean distance transform to ensure stable training (Huttenlocher, Klanderman, & Rucklidge; 2002; Karimi & Salcudean; 2019). Due to its higher computational cost, the HD loss was applied selectively to multi-class experiments, significantly improving contour fidelity in complex organelle reconstructions.

#### Workflow accessibility and reproducibility

All models were implemented in Python (v3.10) using PyTorch and MONAI to ensure compatibility with widely adopted open-source deep learning and bioimaging frameworks. Training and inference were performed on NVIDIA RTX-class GPUs with 8-12 GB of memory, with additional benchmarking conducted on the GRICAD infrastructure using A100 GPUs (32-64 GB). All architectures were optimised for efficient execution on standard laboratory workstations running Windows or Linux. The workflow was developed in accordance with open science principles. A fully archived version of the codebase is available on Zenodo (DOI: https://doi.org/10.5281/zenodo.17630945), including all training scripts, preprocessing pipelines, and experiment configurations. Training datasets and pre-trained model weights are publicly accessible through the BioImage Archive (https://www.ebi.ac.uk/biostudies/bioimages/studies/S-BIAD2407). The modular design of the framework facilitates straightforward integration of additional models or imaging conditions, supporting scalable application to diverse 3D electron microscopy datasets.

## Results

### Performance of baseline architectures for 3D FIB-SEM segmentation

#### Architectural evaluation

We initially evaluated three encoder-decoder architectures with four levels for 3D FIB-SEM segmentation: 3DCNN, 3DSegNet, and 3DVNet (see Figure 3). While all models had similar depth and training configurations, they differed significantly in their upsampling strategies and the use of skip connections, which directly affected segmentation quality (Peng et al., 2023; Zhan et al., 2024).

The 3DCNN featured a straightforward encoder-decoder design without skip connections, resulting in a lightweight architecture with approximately 300,000 parameters. However, the lack of feature reuse between the encoder and decoder limited its ability to recover acceptable boundaries and maintain volumetric continuity On the other hand, 3DSegNet used unparameterised upsampling through pooling indices. This enhanced the model’s capacity to about 10 million parameters and improved spatial reconstruction efficiency. Despite these advantages, boundary delineation proved to be unstable in low-contrast regions. For more details and benchmarked evaluation for 2D see Badrinarayanan et al., 2017.

In contrast, 3DVNet employed learnable transposed convolutions and directly concatenated features between the encoder and decoder stages (Milletari et al., 2016). This design effectively preserved volumetric continuity and fine structural details while remaining computationally efficient, with around 700,000 parameters. All models were trained using the same hyperparameters and loss functions (BCE + Dice) and under identical hardware conditions to ensure a fair comparison (see Table 2).

Although the three architectures share a structurally similar foundation, the differences in feature reuse and upsampling strategies significantly impacted segmentation performance, with 3DVNet consistently providing superior boundary reconstruction and spatial coherence.

#### Region-of-interest versus whole-cell segmentation

To evaluate performance under increasing structural complexity, models were trained and assessed using annotated datasets from the *Phaeodactylum tricornutum* strains Pt1 and Pt18. Strain Pt18, known for its significant morphological variability and diverse chloroplast organisation, served as a rigorous benchmark for model generalisation. We compared two segmentation approaches:

(i) Region-of-Interest (ROI) segmentation, which focuses on localised organelles such as vacuoles and chloroplasts, and (ii) Whole-Cell segmentation, which incorporates background context and inter-organelle relationships.

In the ROI-based segmentation, all models demonstrated high apparent accuracy. The Dice and F_1_ scores for vacuoles were approximately 0.91 for 3DCNN, 0.78 for 3DSegNet, and 0.90 for 3DVNet. For chloroplast segmentation, the Dice values were around 0.97 for 3DCNN, 0.89 for 3DSegNet, and 0.97 for 3DVNet (see Figure 4A-D). These results indicate that ROI segmentation is a relatively manageable task, even for superficial architectures.

**Figure 4.**
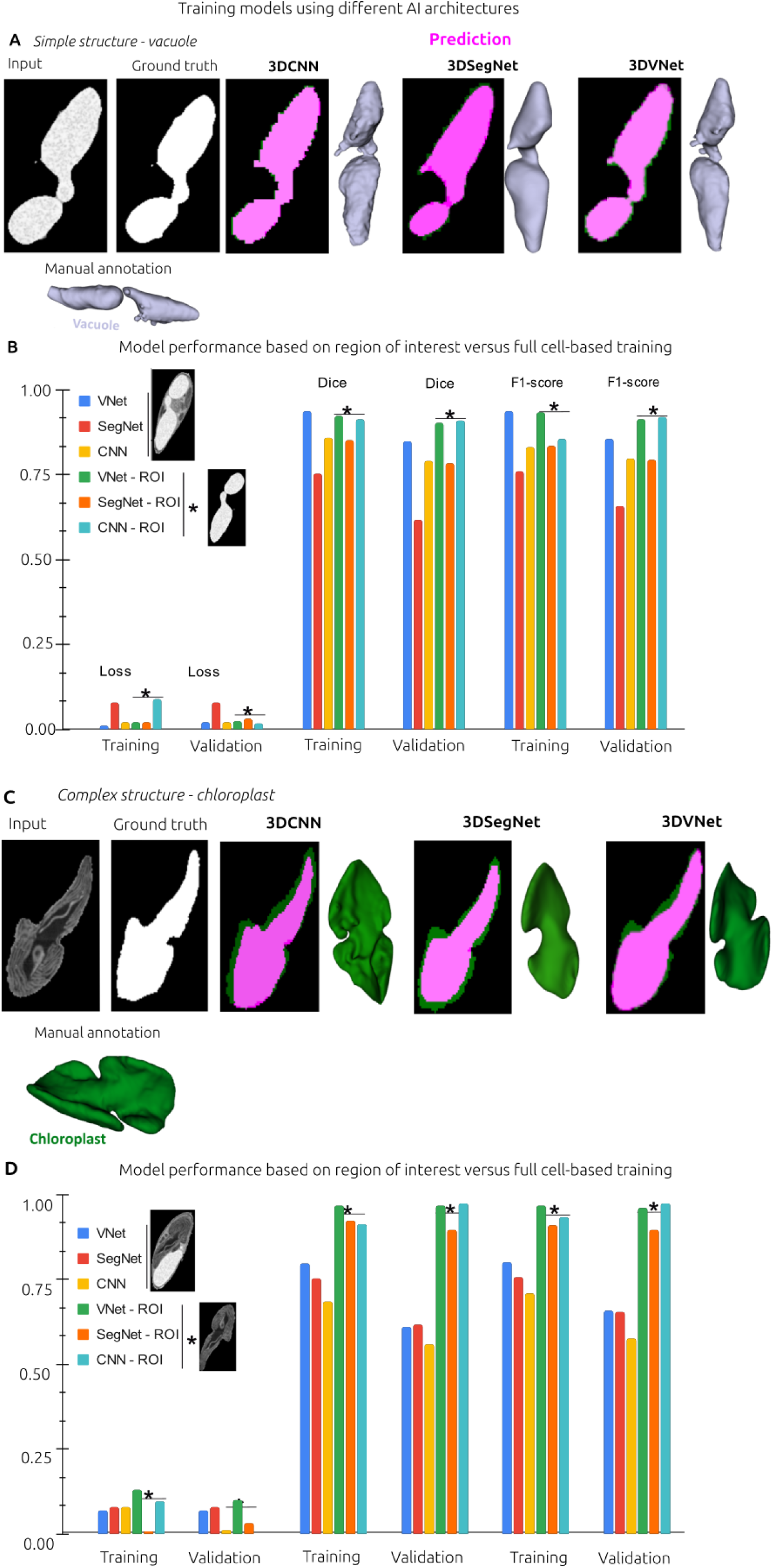
Benchmarking CNN-based architectures and training strategies for 3D FIB-SEM segmentation. (A, C) Representative segmentation results for simple structures (vacuoles) and complex structures (chloroplasts) obtained using 3DCNN, 3DSegNet, and 3DVNet architectures. Ground-truth annotations (white) are compared with predicted segmentations (pink), with corresponding 3D reconstructions shown alongside manual annotations. (B, D) Quantitative comparison of models trained using region-of-interest (ROI)-based learning versus whole-cell training. Bars represent Dice coefficient, F1-score, and loss for training and validation sets. Asterisks (*) indicate improved performance achieved through ROI-based training, particularly for fine structural details.

In contrast, whole-cell segmentation led to a significant decrease in performance across all models (see Figure 4B, D). The Dice scores for vacuoles were 0.85 for 3DVNet, 0.79 for 3DCNN, and 0.62 for 3DSegNet, while chloroplast Dice scores ranged from 0.60 to 0.80 across architectures. When applying a minimum acceptable Dice threshold of 0.70, 3DVNet consistently achieved the best balance between boundary precision and contextual stability. On the other hand, 3DCNN exhibited overfitting, whereas 3DSegNet displayed unstable convergence.

These findings underscore that, although ROI segmentation may overestimate model performance, whole-cell segmentation highlights significant differences in architectural robustness. As structural complexity increased, 3DVNet emerged as the most reliable architecture, justifying its selection for next experiments.

#### Effect of extended training and dataset variability

Extending 3DVNet training to 1,200 epochs further improved the recovery of fine structural details without increasing computational cost (see Figure 5A), achieving performances comparable to that of previously reported deeper architectures (Wang et al. 2022). When trained on the morphologically simpler Pt1 dataset, convergence was reached within ∼200 epochs (Dice ≈ 0.90; F_1_ ≈ 0.86). In contrast, 3DCNN and 3DSegNet required longer optimisation. They exhibited greater instability (see SFigure 1) These results highlight 3DVNet’s scalability across datasets with varying morphological complexity and support its suitability as a baseline architecture.

**Figure 5.**
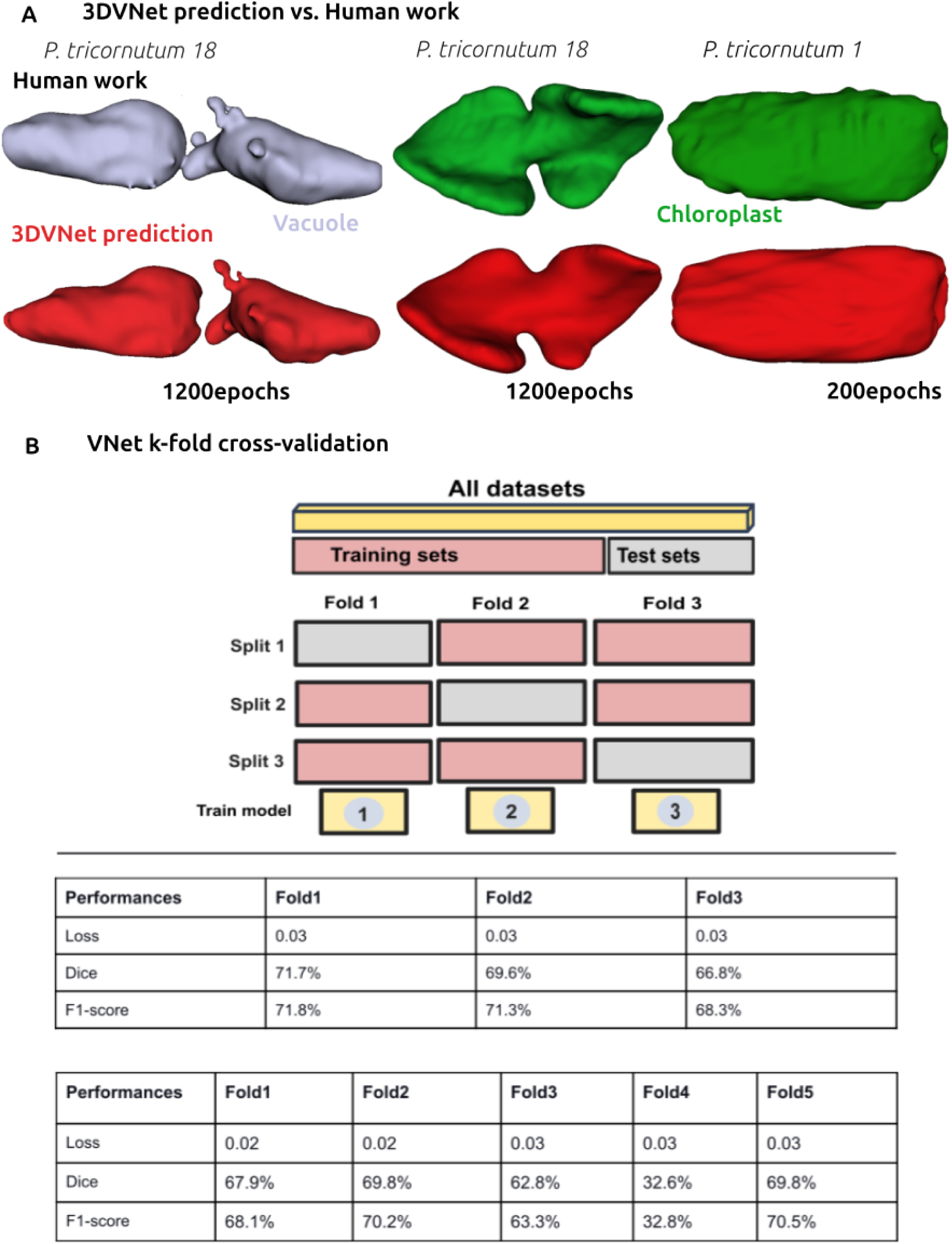
Evaluation of 3DVNet predictions and k-fold cross-validation performance. (A) Comparison between manual segmentations (“Human work”) and 3DVNet predictions for vacuoles and chloroplasts in *Phaeodactylum tricornutum* strains Pt18 and Pt1. Manual annotations are shown in grey and green, while model predictions (red) are shown after 1200 and 200 training epochs, respectively. (B) K-fold cross-validation of the 3DVNet model across 32 annotated FIB-SEM volumes. Schematics illustrate both 3-fold and 5-fold configurations, and tables summarise loss, Dice coefficient, and F1-score for each fold. All models were trained using identical preprocessing and hyperparameters (see Table 2).

### Cross-validation and model robustness

To assess generalization, we combined the Pt1 and Pt18 datasets into a composite collection of 32 annotated volumes that contains a diverse range of morphotypes. We applied both three-fold and five-fold cross-validation to the 3DVNet (see Figure 5B). The three-fold cross-validation produced stable convergence, with loss variability ranging from 0.02 to 0.03 and average Dice/F_1_ scores exceeding 0.70 across the folds. In contrast, the five-fold cross-validation demonstrated higher variance and reduced performance in morphologically extreme subsets, with Dice scores dropping to approximately 0.33. These results suggest that moderate cross-validation methods strike the best balance between the volume of training data and the diversity of the dataset for small, heterogeneous 3D electron microscopy datasets.

### Benchmarking against MONAI architectures

We benchmarked 3DVNet against two widely used MONAI architectures (Cardoso et al., 2022), BasicUNet and DynUNet, using the same preprocessing, training protocols, and evaluation metrics (see Figure 6). The training datasets included combined volumes from Pt1 and Pt18 (a total of 32 stacks), and we used manually curated vacuole and chloroplast annotations as reference masks. Our qualitative comparisons (Figure 6A) and quantitative metrics (Figure 6B) revealed that both VNet and DynUNet achieved consistent segmentation quality, while BasicUNet demonstrated reduced robustness. This benchmarking established a reliable performance baseline and highlighted potential architectures for transfer learning and hybridisation.

**Figure 6.**
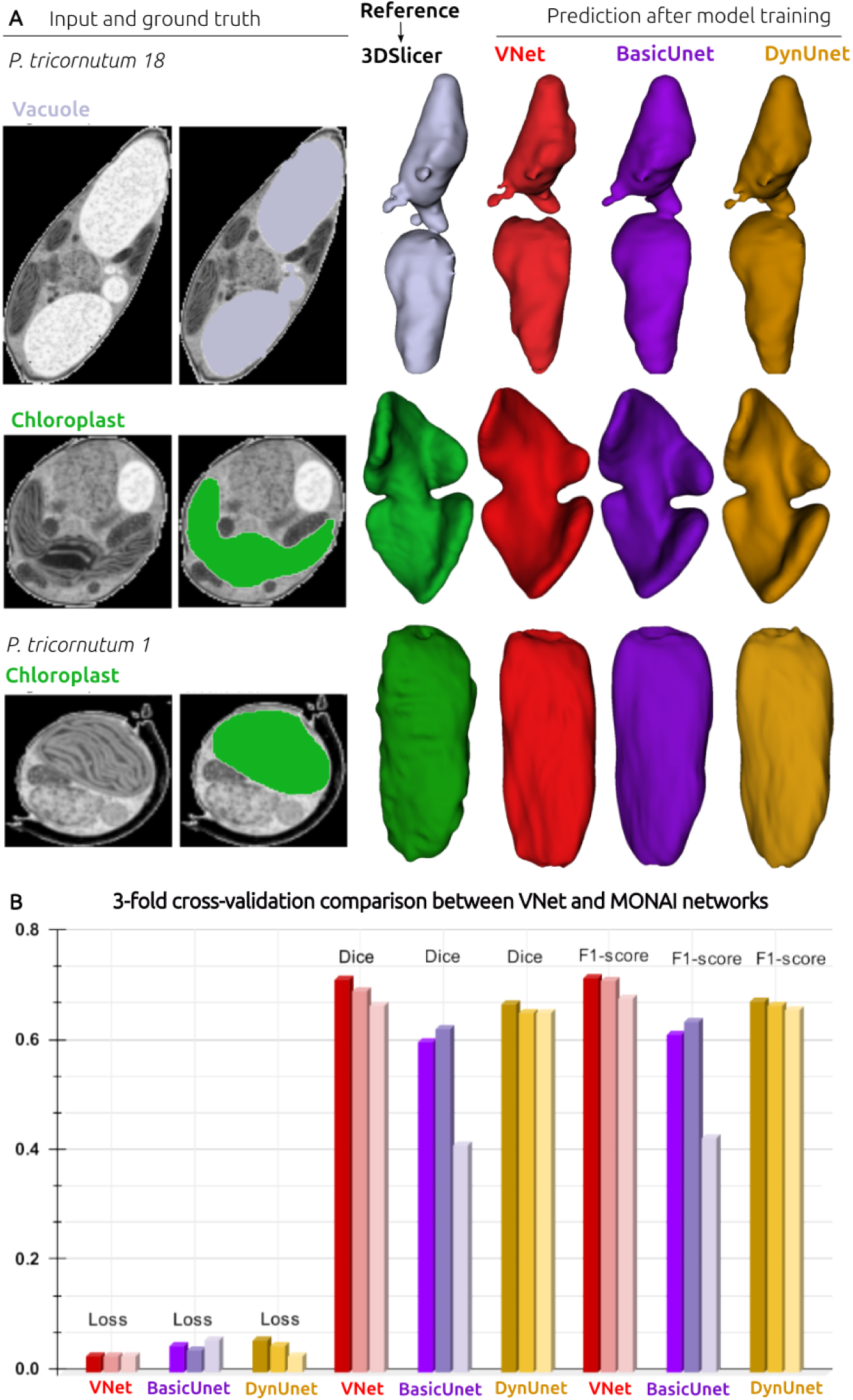
Comparative benchmarking of VNet, BasicUNet, and DynUNet architectures on 3D FIB-SEM datasets. (A) Examples of input FIB-SEM images, manual annotations, and 3D reconstructions for Pt18 and Pt1. Ground-truth segmentations for vacuoles (light purple) and chloroplasts (green) were generated using 3D Slicer. Model predictions are shown for VNet (red), BasicUNet (purple), and DynUNet (gold), all trained under identical conditions. (B) Quantitative comparison of validation metrics, including loss, Dice coefficient, and F1-score, for each architecture. All results correspond to models trained on the combined Pt18 and Pt1 datasets using identical preprocessing and hyperparameters.

### Transfer learning and hybrid frameworks

#### Transfer learning on simple structures

All models were pretrained on the combined Pt1 and Pt18 datasets and thereafter fine-tuned on individual strains. For vacuole segmentation, pretrained VNet and DynUNet achieved comparable performance, while BasicUNet demonstrated limited transferability (Figure 7A). Hybrid architectures combining DynUNet + VNet and Swin-Net + VNet were also evaluated. While DynUNet + VNet did not outperform DynUNet alone, the Swin-Net + VNet hybrid achieved the highest proportion of qualitatively “good” outcomes across 39 unseen samples (Figure 7B).

**Figure 7.**
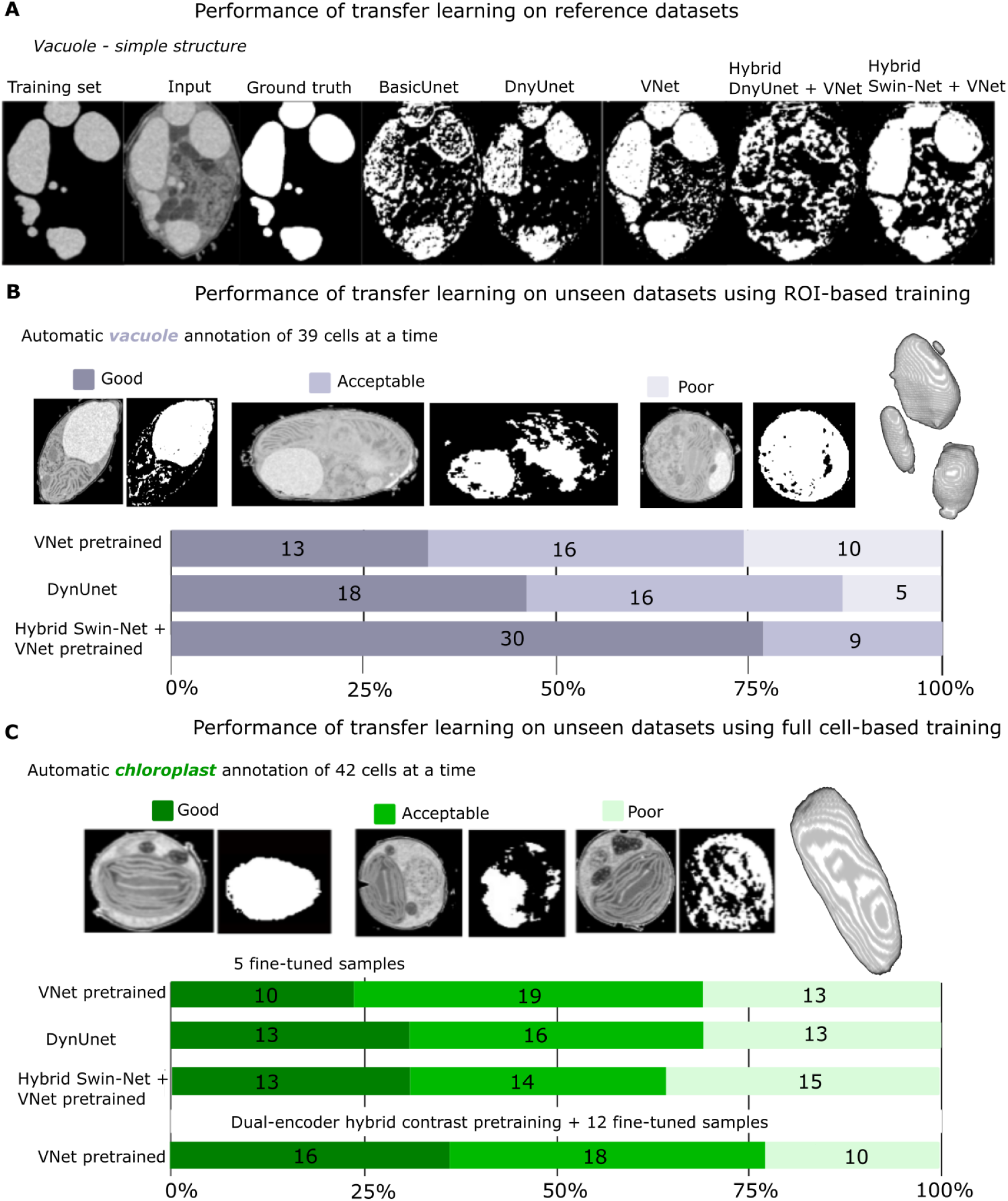
Transfer learning and contrast-aware hybrid segmentation of 3D FIB-SEM datasets. (A) Performance of transfer learning on reference datasets illustrated using representative vacuole segmentations. Shown are input FIB-SEM images, manual ground-truth annotations, and predictions from pretrained models (BasicUNet, DynUNet, VNet) and hybrid configurations (DynUNet + VNet, Swin-Net + VNet). (B) Transfer learning performance on unseen datasets was evaluated using ROI-based training. Automatic vacuole segmentation of 39 cells was qualitatively categorised as Good, Acceptable, or Poor. Bar charts summarise the distributions of outcomes for each pretrained or hybrid model. (C) Transfer learning performance on unseen datasets using whole-cell training for chloroplast segmentation. Fine-tuning was performed with either five or twelve annotated samples. Results for a dual-encoder contrast-aware VNet, trained separately on low- and high-contrast datasets, are shown. Bar charts summarise qualitative performance across 42 cells for each condition.

#### Transfer learning on complex morphologies

For chloroplast segmentation, both pretrained and hybrid models were fine-tuned using either 5 or 12 annotated samples. The increase in supervision significantly improved performance across all models. While the combination of Swin-Net and VNet modestly improved segmentation continuity, a contrast-aware dual-encoder VNet trained separately on high- and low-contrast subsets achieved the highest robustness. This model recorded Dice scores of approximately 0.82 for low contrast and around 0.88 for high contrast (see Figure 7C). These results demonstrate that contrast-aware learning significantly enhances generalisation across diverse FIB-SEM acquisitions. The results indicate that contrast-aware learning significantly improves generalisation across various FIB-SEM acquisitions. This is particularly evident in organelle segmentation in volumetric electron microscopy, as discussed by Heinrich et al. (2021) and Wei et al. (2020). In summary, these thorough experiments highlight the potential of pretrained architectures to enable quick adaptation to new volumetric datasets.

#### Loss functions and multi-class segmentation

Extending segmentation techniques to multi-class scenarios has introduced challenges related to spatial overlap and grayscale similarity among organelles. Models trained using Binary Cross-Entropy (BCE) loss maintained overall morphology but showed softened boundaries and organelle merging (see Figure 8A and 8B). By incorporating the Hausdorff Distance (HD) loss, we observed significant improvements in boundary delineation and a reduction in inter-organelle overlap (see Figure 8C). Both DynUNet and VNet benefited from optimisation using the HD loss. At the same time, the hybrid model combining Swin-Net and VNet further enhanced contour coherence through multi-scale contextual encoding. Although using HD loss increased computational cost, it yielded segmentations with superior geometric fidelity and closer alignment with expert annotations, which is collaborated by Ma et al. (2021) and Kalimi et al. (2019). These findings demonstrate that while BCE loss is appropriate for lightweight benchmarking and single-class segmentation, boundary-aware losses are crucial for achieving biologically realistic reconstructions of multiple organelles.

**Figure 8.**
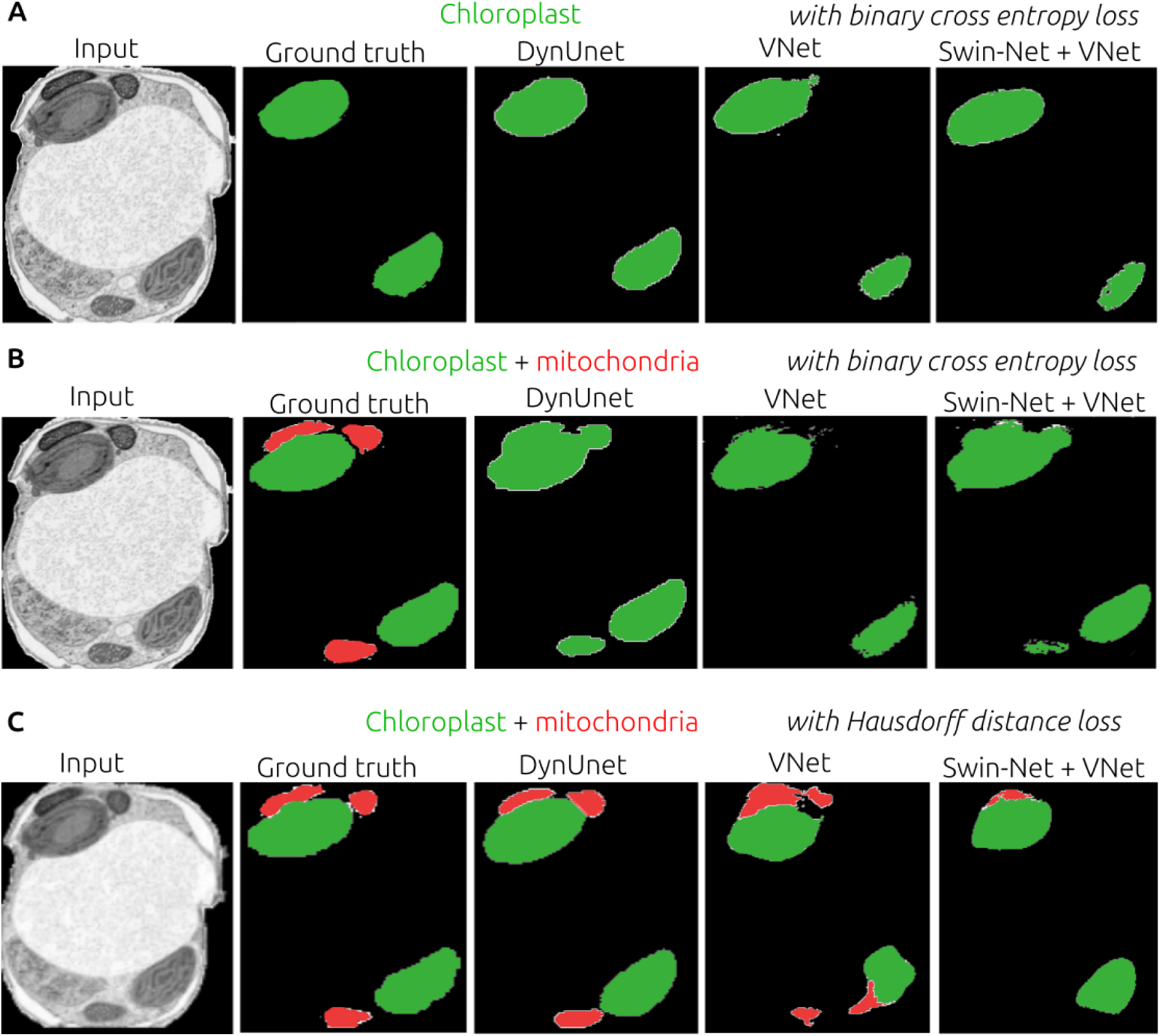
Comparison of loss functions for single- and multi-class segmentation. (A) Single-class chloroplast segmentation using models trained with Binary Cross-Entropy (BCE) loss. (B) Multi-class segmentation of chloroplasts and mitochondria using BCE loss. (C) Multi-class segmentation using Hausdorff Distance (HD) loss. Shown are input FIB-SEM slices, ground-truth annotations, and predictions from DynUNet, VNet, and the Swin-Net + VNet hybrid. Chloroplasts are shown in green and mitochondria in red.

## Conclusion and remarks

This study presents a systematic framework for evaluating and optimising deep learning architectures for the 3D segmentation of subcellular structures in FIB-SEM datasets. By benchmarking both conventional and advanced models under controlled conditions, we demonstrate that segmentation performance is influenced by network architecture, data diversity, and loss-function design. By benchmarking both conventional and advanced models under controlled conditions, we demonstrate how the combined influence of network architecture, data diversity, and loss function design affects segmentation performance. Among the approaches evaluated, VNet proved to be the most effective, showing an optimal balance between computational efficiency and volumetric accuracy. Additionally, DynUNet displayed particular strengths in multi-class segmentation tasks that require precise boundary delineation. We also show that pretrained models can be efficiently adapted to new datasets with minimal annotation through transfer learning and contrast-aware hybrid strategies. This adaptability enables scalable applications across morphologically diverse microalgal species. The incorporation of boundary-aware loss functions significantly improved the separation of closely associated organelles, such as chloroplasts and mitochondria. This highlights the importance of combining voxel-level accuracy with geometric constraints for achieving biologically realistic segmentation.

However, our findings reveal significant practical limitations. Training on regions of interest can increase apparent performance compared to whole-cell inference, but background context, rare morphotypes, and contrast variability expose robustness limits. Furthermore, achieving broader generalisation across different species, preparation protocols, and extreme contrast conditions will require extended training datasets and targeted fine-tuning. Future developments integrating domain adaptation strategies, organelle-specific encoders, and user-friendly graphical user interfaces (GUIs) are expected to enhance transferability , annotation and facilitate performance in high-throughput comparative analysis of ultrastructural phenotyping.

## Data availability

All data supporting the findings of this study are available within the article and its Supplementary Information. Raw FIB-SEM image stacks, ground truth annotations, and pretrained model weights are deposited in the BioImage Archive under the accession number S-BIAD2407.

## Code availability

The code used for training, benchmarking, and evaluating the transfer learning of neural network architectures is available on [GitHub](https://github.com/cuwizeye/AI-assisted-learning) and permanently archived on [Zenodo](https://zenodo.org/records/17630945). The repository includes command-line scripts for data preprocessing, model training, and performance evaluation, all of which are built using **PyTorch** and **MONAI.** The framework is designed for reproducibility and open adaptation; however, it currently operates via command-line execution. A graphical user interface (GUI) is in development for future releases.

## Acknowledgements

We acknowledge funding from the European Research Council (ERC) for the Chloro-Mito project (grant no. 833184 to G. F., S. F. and J. A.) and for the PlanktON project (grant no. 101099192 to C. U., G. F. and P. A.) from the European Innovation Council (EIC). P.A. also acknowledges support from the European Union’s Horizon 2020 research and innovation programme via the Marie Skłodowska-Curie grant agreement no. 101066400 (PhotoLINK). Computational resources were provided by the GRICAD cluster infrastructure (https://gricad.univ-grenoble-alpes.fr), supported by the Grenoble research community. We thank **Mondher Chekki** for granting access to additional computational resources, which enabled large-scale model training and validation. Special acknowledgement goes to **Leandro Farias Estrozi** for providing feedback on an earlier draft.

## Supporting informations

### 1. Architectural flow summary between different neural network

**STable 1:**
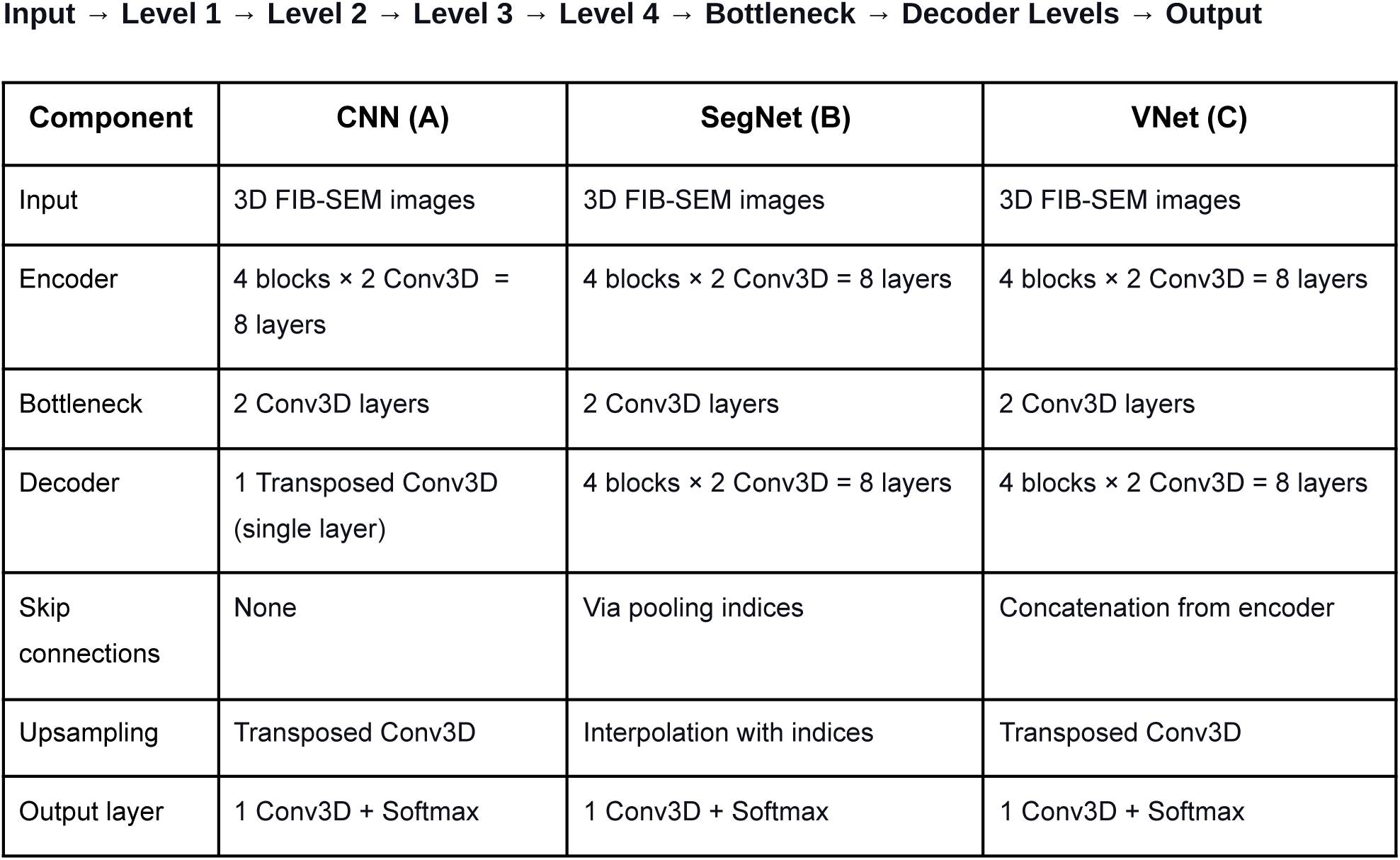
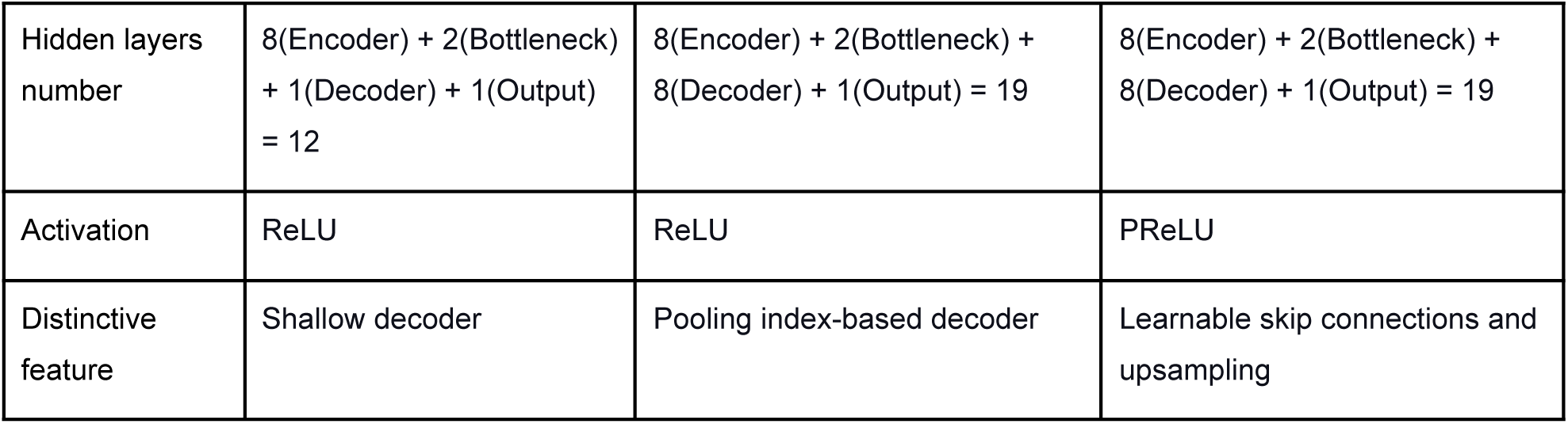
The main components of the neural network architecture are shown for the CNN (A), SegNet (B) and VNet (C) architectures.

### 2. Neural network architecture encoder-decoder blocks construction

**STable 2:**
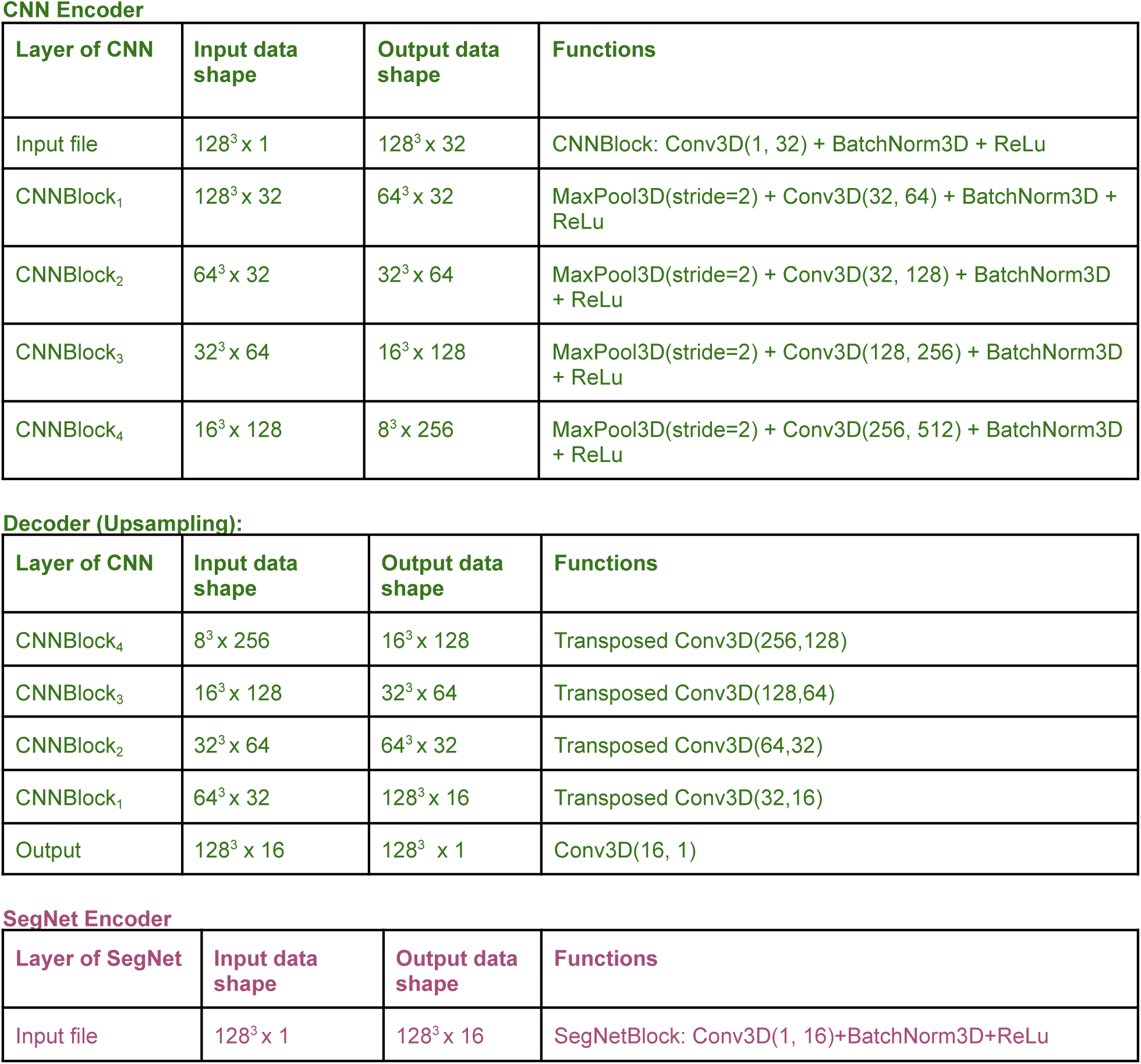
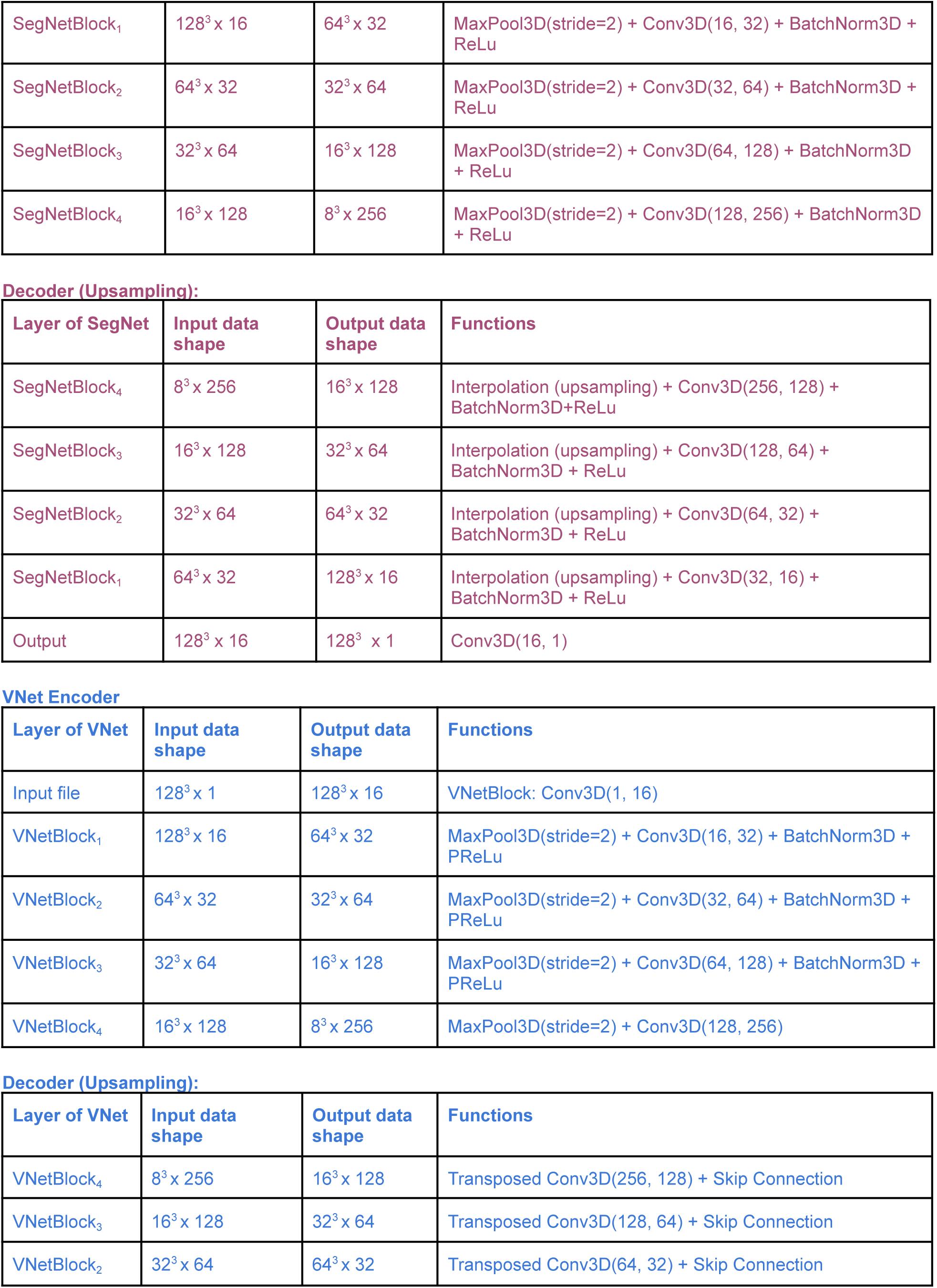
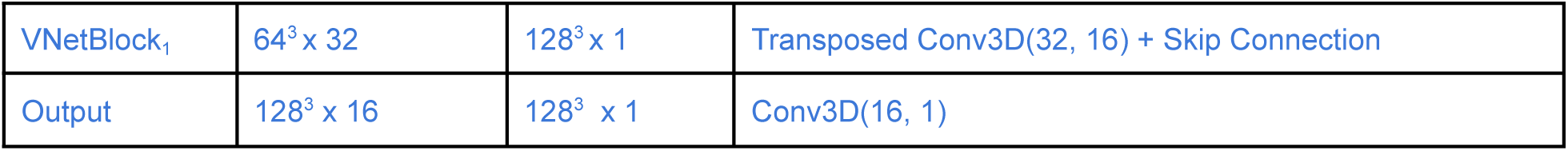
*Details of the used neural network architecture*. Shows input/output shape of the data and the operations applied at each layer, including the specifics of the encoder and decoder blocks.

**SFigure1.**
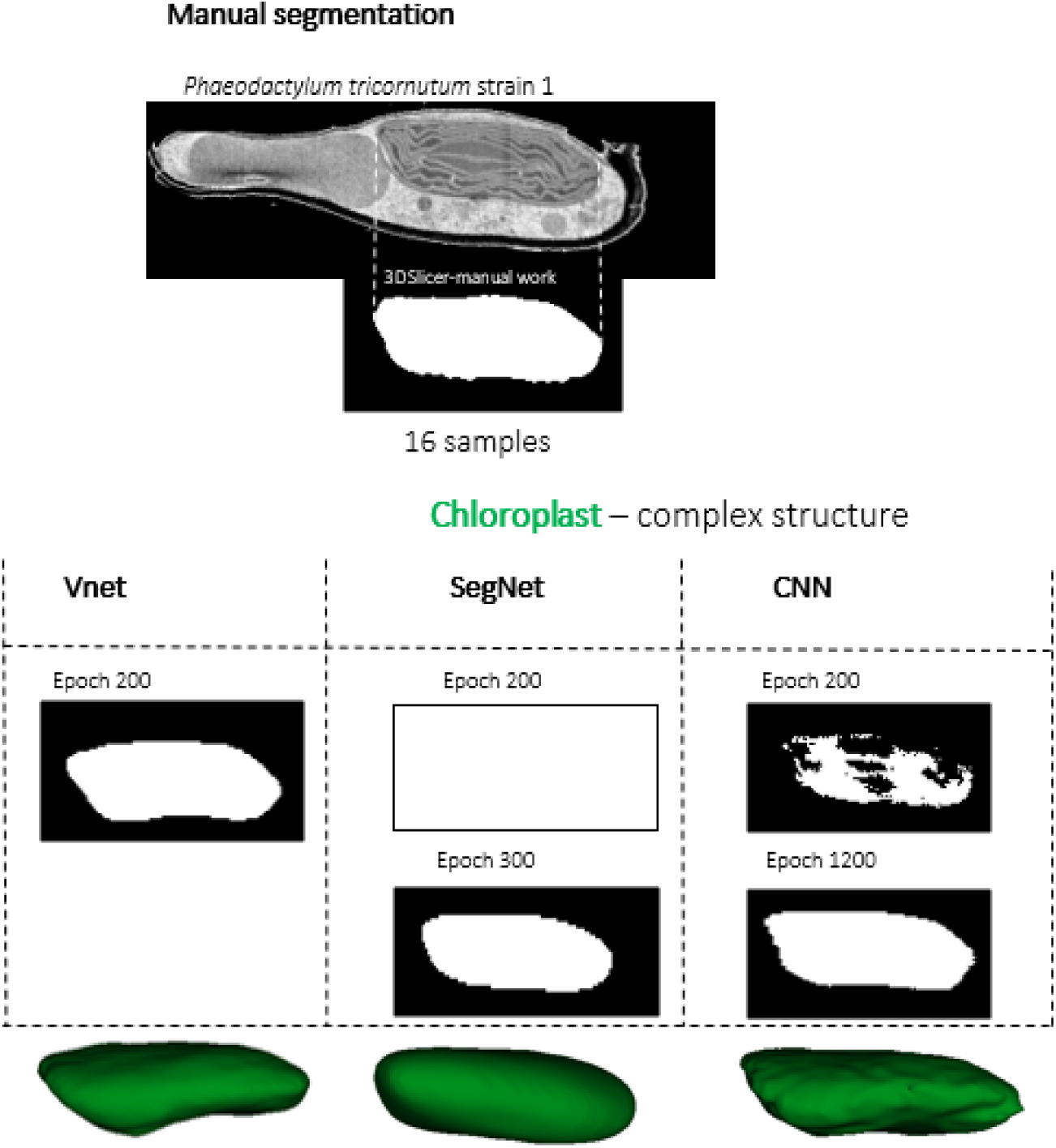
Comparative segmentation performance of VNet, SegNet, and CNN on complex chloroplast structures. Manual annotations of the *Phaeodactylum tricornutum* strain Pt1 (top) were generated using 3D Slicer on 16 annotated samples, which serve as the ground truth for model evaluation. The segmentation results for chloroplasts are presented for three different architectures: 3DVNet, 3DSegNet, and 3DCNN, each trained under identical conditions across varying numbers of epochs (ranging from 200 to 1200).

